# *Komagataella phaffii* encodes two functional Pho4 transcription factors

**DOI:** 10.64898/2026.02.17.706399

**Authors:** Marcel Albacar, Asier González, Rui Wang, Antonio Casamayor, Joaquín Ariño

## Abstract

The transcription factor Pho4 is crucial for the response to phosphate starvation in many fungi, and it has been linked to tolerance to alkalinization of the medium and to pathogenicity. It is widely accepted that it is encoded by a single gene. However, the industrially relevant yeast *Komagataella phaffii* might contain two Pho4-encoding genes (*PAS_chr1-1_0265* and *PAS_chr2-1_0177*, designated here *PHO4(A)* and *PHO4(B*), respectively), which have never been functionally characterized. The phenotypic analysis of single and double mutants suggests that Pho4(B) plays a major role in the adaptation to Pi scarcity. While single mutants exhibited limited and non-overlapping phenotypic defects, the *pho4(A) pho4(B)* strain was sensitive to multiple types of stress, including phosphate starvation and alkaline pH. Transcriptomic analysis confirms that Pho4(B) is crucial for the transcriptional response to phosphate starvation, including induction of typical gene markers (*PHO5*, *PHO89*, *VTC1*, etc.). However, by using a GFP reporter we found that *PHO4(A)* also participates in the induction of *PHO89* under high pH stress. Expression of both *PHO4(A)* and *PHO4(B)* in *S. cerevisiae* complemented the *pho4* mutation under phosphate limitation by restoring growth, expression of the Pho84 transporter and secreted phosphatase activity. These results indicate that both transcription factors display partially overlapping functions, responding differently to diverse stimuli, and that together they constitute a key component in the adaptation to a variety of stresses. Therefore, *K. phaffii* is an exceptional example among fungi that encodes two Pho4 functional transcription factors.

## INTRODUCTION

Phosphate is a fundamental nutrient required for key cellular processes, including energy metabolism, nucleic acid synthesis, membrane formation, and signal transduction. Environmental phosphate availability can fluctuate widely, especially in the natural and industrial niches yeast occupy. The ability to sense and respond to variation of phosphate levels allows yeasts to maintain growth and viability by activating phosphate acquisition systems, mobilizing internal phosphate reserves, and adjusting metabolic pathways to conserve this resource (Austin and Mayer 2020; Mouillon and Persson 2006).

Adaptation to inorganic phosphate (Pi) scarcity has been studied in detail in the yeast *Saccharomyces cerevisiae.* Under conditions of phosphate repletion, inositol phosphates (possibly 1,5-IP_8_) inactivate the cyclin-dependent kinase inhibitor Pho81 through its SPX domain (Chabert et al. 2023; Lee et al. 2007) which leads to the activation of the cyclin-dependent kinase (CDK) complex Pho85-Pho80 (Ogawa et al. 1995; Yoshida et al. 1989). This complex phosphorylates the crucial helix-loop-helix (HLH) Pho4 transcription factor (TF) at several Ser-Pro (SP) motifs, leading to its inactivation and cytosolic location by means of the exportin Msn5 (Kaffman et al. 1994, 1998a; Komeili and O’Shea 1999; O’Neill et al. 1996; Fung et al. 2025). In contrast, during Pi starvation, the inhibition of Pho85-Pho80 kinase by Pho81 leads to the dephosphorylation of Pho4 and its entry to the nucleus (Kaffman et al. 1998b), thus resulting in the expression of ≈20 genes required for survival under Pi limitation (the so called, PHO regulon), such as those encoding high affinity Pi transporters (*PHO84* and *PHO89*), or various secreted phosphatases (*PHO5* and *PHO11*). In *S. cerevisiae*, effective gene induction requires the cooperative binding to the promoters of another transcription factor, Pho2 (Bas2), which results in increased selectivity of the response to phosphate starvation (Zhou and O’Shea 2011). Although this response mechanism is well conserved, some differences are found among fungi with regard the requirement for Pho2 cooperation. Thus, Pho2 exists in *Candida albicans* and *Candida glabrata,* but its absence does not affect the ability to grow on Pi limitation conditions (He et al. 2017; Kerwin and Wykoff 2009), whereas no Pho2 orthologs have been identified in *Cryptococcus neoformans* (Gomes-Vieira et al. 2018). An extreme case is the fission yeast *Schizosaccharomyces pombe*, in which the response to phosphate starvation is mediated by a transcription factor structurally different from Pho4, the Zn_2_Cys_6_ binuclear cluster-containing Pho7 (Carter-O’Connell et al. 2012; Schwer et al. 2017). In all these cases, such variations coincide with an expanded range of genes whose expression depends on Pho4 (Lev and Djordjevic 2018).

The adaptive responses to alkalinization and to phosphate starvation have many points in common. In *S. cerevisiae*, Pho4 also migrates to the nucleus upon alkalinization (Serra-Cardona et al. 2015) and it is required for the induction of *PHO84* and *PHO12* (Serrano et al. 2002). Concordantly, deletion of *PHO2*, *PHO4* or *PHO81* results in strong sensitivity even to moderate alkalinization (Serrano et al. 2004; Ariño 2010). Similarly, mutation of *PHO4* in *C. albicans* (Ikeh et al. 2016) and in *C. neoformans* (Lev et al. 2017) results in growth defects not only under Pi limitation but also at alkaline pH. Notably, lack of Pho4 attenuates the virulence of these two fungal pathogens (Urrialde et al. 2016; Ikeh et al. 2016; Lev et al. 2017).

*Komagataella phaffii* (formerly *Pichia pastoris*) is a methylotrophic yeast well known for its use in heterologous protein production. Its strengths include the capability to grow at high cell density, to carry out eukaryotic post-translational modifications, efficient secretion of the produced proteins into the environment, and the availability of appropriate genetic modification tools. (Wu et al. 2023; Barone et al. 2023; Love et al. 2018; Vijayakumar and Venkataraman 2024). Surprisingly, despite its importance, our knowledge about how *K. phaffii* adapts to Pi scarcity is virtually null and the first hint about how this yeast remodels gene expression under Pi limitation has been reported only recently (Ishtuganova et al. 2024). Our laboratory is interested in developing novel platforms for protein expression based on the use of promoters that are regulated by mild alkalinization both in *S. cerevisiae* and in *K. phaffii* (Albacar et al. 2023, 2024; Zekhnini et al. 2025) and we have recently charted in *K. phaffii* the contribution of the transcription factors Crz1 and Rim101, which are key components of two signaling pathways crucial for adaptation to alkalinization of the environment (Albacar et al. 2025). Given the functional links between Pi starvation and alkaline pH response, we initially considered investigating the possible role of Pho4 for adaptation to high pH. The signaling machinery for Pho4 regulation seems reasonably well conserved in *K. phaffii* when compared with other fungi, such as *S. cerevisiae* (Figure 1A). However, while Pho4 is found in fungi as a single gene, our search of the *K. phaffii* genome suggested that it might contain two genes, *PAS_chr1-1_0265* and *PAS_chr2-1_0177*, encoding possible Pho4 proteins of 380 and 365 amino acids, respectively. Notably, these proteins displayed rather low identity among them as well as to other well-characterized fungal Pho4 proteins (see Results). This raised a key question: Does *K. phaffii* encode two Pho4 transcription factors? To address this issue, we created single and double mutants for these genes and used them to carry out detailed phenotypic analyses, and transcriptomic profiling in response to phosphate starvation, as well as complementation experiments in *S. cerevisiae*. Our data supports the notion that, although the functions of *PAS_chr1-1_0265* and *PAS_chr1-1_0177* (named here *PHO4(A)* and *PHO4(B)*, respectively) do not fully overlap, they both behave as Pho4 TF in *K. phaffii*. Adaptation to low Pi conditions relies largely on Pho4(B). While cells lacking individual TFs are affected by certain specific stress conditions, the double mutation yields cells sensitive to multiple stresses, indicating that, in combination, these proteins are key components in the stress response. In addition, we show that both proteins can replace Pho4 in *S. cerevisiae*. Therefore, *K. phaffii* exceptionally encodes two functional Pho4 transcription factors.

**Figure 1.**
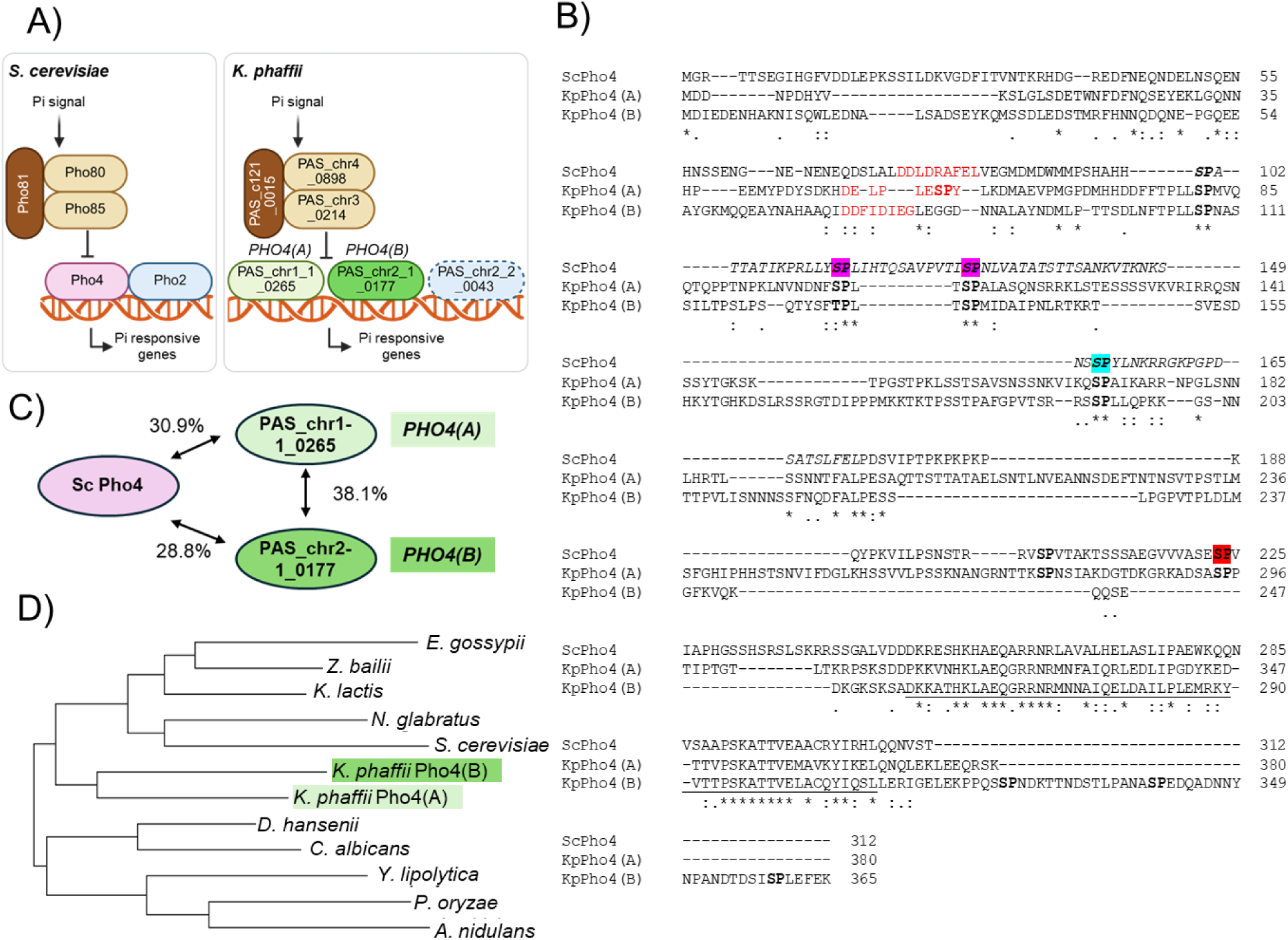
Identification of two genes encoding putative Pho4 transcription factors in *K. phaffii*. A) Downstream components of the regulon involved in the response to Pi starvation in *S. cerevisiae* (left) and the putative components of the system in *K. phaffii*. B) Clustal Ω alignment of *S. cerevisiae* Pho4 (Uniprot P07270), and *K. phaffii PAS_chr1-1_0265* (KpPho4(A)) and *PAS_chr2-1_0177* (KpPho4(B)) protein products. The bHLH DNA-binding domains are underlined. Phosphorylatable dipeptides SP and TP are highlighted in bold (the alignment was manually edited to align better the known Pho85 recognition motifs). Residues whose phosphorylation is required for ScPho4 nuclear export (sites 2 and 3) are denoted in magenta, and the ones required for inhibition of nuclear import (site 4) and Pho2 interaction (site 6) in blue and red, respectively. The region in *S. cerevisiae* Pho4 involved in nuclear import and export appears in italics. The known *S. cerevisiae* 9 aaTAD (McAndrew et al. 1998; Piskacek et al. 2007) and the predicted equivalent regions for *K. phaffii* proteins are denoted in red font. C) Amino acid identity relationships between *S. cerevisiae* Pho4, and *K. phaffii* Pho4(A) and Pho4(B) proteins. D) Phylogenetic analysis of Pho4 proteins from 11 fungal species. Note that a single protein is identified in all cases except for *K. phaffii*.

## RESULTS

### Identification of *PHO4* candidates in *K. phaffii*

BLASTp analysis of the NCBI reference proteome database using *S. cerevisiae* Pho4 (Uniprot entry P07270) as bait revealed two possible hits in the *K. phaffii* GS115 proteome (Supplemental_Table_S1). The first hit (E-value 4.0E-10) corresponded to the hypothetical protein of 380 amino acids encoded by *PAS_chr1-1_0265*, named here Pho4(A), and the second (E-value 3.0E-7) to the hypothetical protein of 365 residues encoded by *PAS_chr2-1_0177*, named here Pho4(B). However, the similarity was circumscribed to the predicted basic helix–loop–helix (bHLH) DNA binding domain (Figure 1B). In fact, the overall identity in comparison with *S. cerevisiae* Pho4 was rather low: 30.9% for Pho4(A) and 28.8% for Pho4(B) (Figure 1C), and both *K. phaffii* proteins were only distantly related to each other (38.1% identity). As for ScPho4, and apart from the bHLH-like region, AlphaFold predicts an intrinsically disordered structure for KpPho4(A) (Supplemental_Fig_S1). This is also the case for KpPho4(B), except that a α–helical region is predicted for the first 42 residues. As mentioned in the Introduction, the existence of two genes encoding different Pho4 proteins is an anomaly because, when present, only one putative Pho4-encoding gene is found (Figure 1D and Supplemental_Table_S1). Nevertheless, close examination revealed in both proteins several putative Pho85 SP phosphorylation sites, as well as a possible 9 amino acids transcriptional activation domain (9aaTAD). These elements are reminiscent of those that proved to be relevant in *S. cerevisiae* for regulation of Pho4 activity and subcellular localization (see Figure 1B and the Introduction section). This suggested that, despite their distant relationship in primary sequence, both proteins could function as *bona fide* Pho4 transcription factors. We therefore proceeded with their functional characterization.

### Phenotypic analysis of single and double Pho4 *K. phaffii* mutants reveals both specialized and redundant functions

In diverse fungi lack of Pho4 function leads not only to inability to transcriptionally respond to phosphate starvation but also to increased sensitivity to diverse forms of stress. Therefore, we exposed strains lacking Pho4(A), Pho4(B) or both proteins to a set of stress conditions and evaluated the resulting phenotypes. As shown in Figure 2A, growth of cells mutated for *PHO4(A)* did not differ from the wild-type strain under Pi limitation, whereas *pho4(B)* mutants exhibited a slight growth defect in both liquid and plate cultures that was aggravated in the double mutant. Cells lacking either Pho4(A) or Pho4(B) exhibited normal tolerance to alkaline pH (Figure 2B) or high concentrations of Ca^2+^, whereas the absence of both transcription factors caused a dramatic loss of tolerance to high pH and a noticeable growth defect in the presence of calcium. Notably, *pho4(A)* cells, but not *pho4(B)* mutants, were sensitive to Mn^2+^ cations, and this phenotype was aggravated in the double mutant. Neither single nor double mutants displayed altered tolerance to NaCl, although the double mutant grew slowly above 0.4 M NaCl (Figure 2B). Remarkably, these cells did not show increased sensitivity to Li^+^ cations, a more toxic analog of Na^+^, suggesting that the growth defect in the presence of high concentrations of NaCl could be derived from the associated osmotic shock. In agreement with this hypothesis, *pho4(A) pho4(B)* cells grew poorly in the presence of high amounts of sorbitol (Figure 2B). The *pho4(A)* mutant, but not the *pho4(B)*, was slightly sensitive to Zn^2+^ ions, and the effect was intensified by the additional mutation of *PHO4(B)*. The double mutants were also unable to grow at 37 °C, whereas cells lacking Pho4(A) or Pho4(B) grew similarly to the wild-type strain. Finally, *pho4(A)* mutants behaved as the wild-type strain in the presence of the anionic detergent sodium dodecyl sulfate (SDS), whereas mutation of *PHO4(B)* resulted in increased sensitivity that was further exacerbated by the additional deletion of *PHO4(A)* (Figure 2B). A summary of these results is presented in Figure 2C.

**Figure 2.**
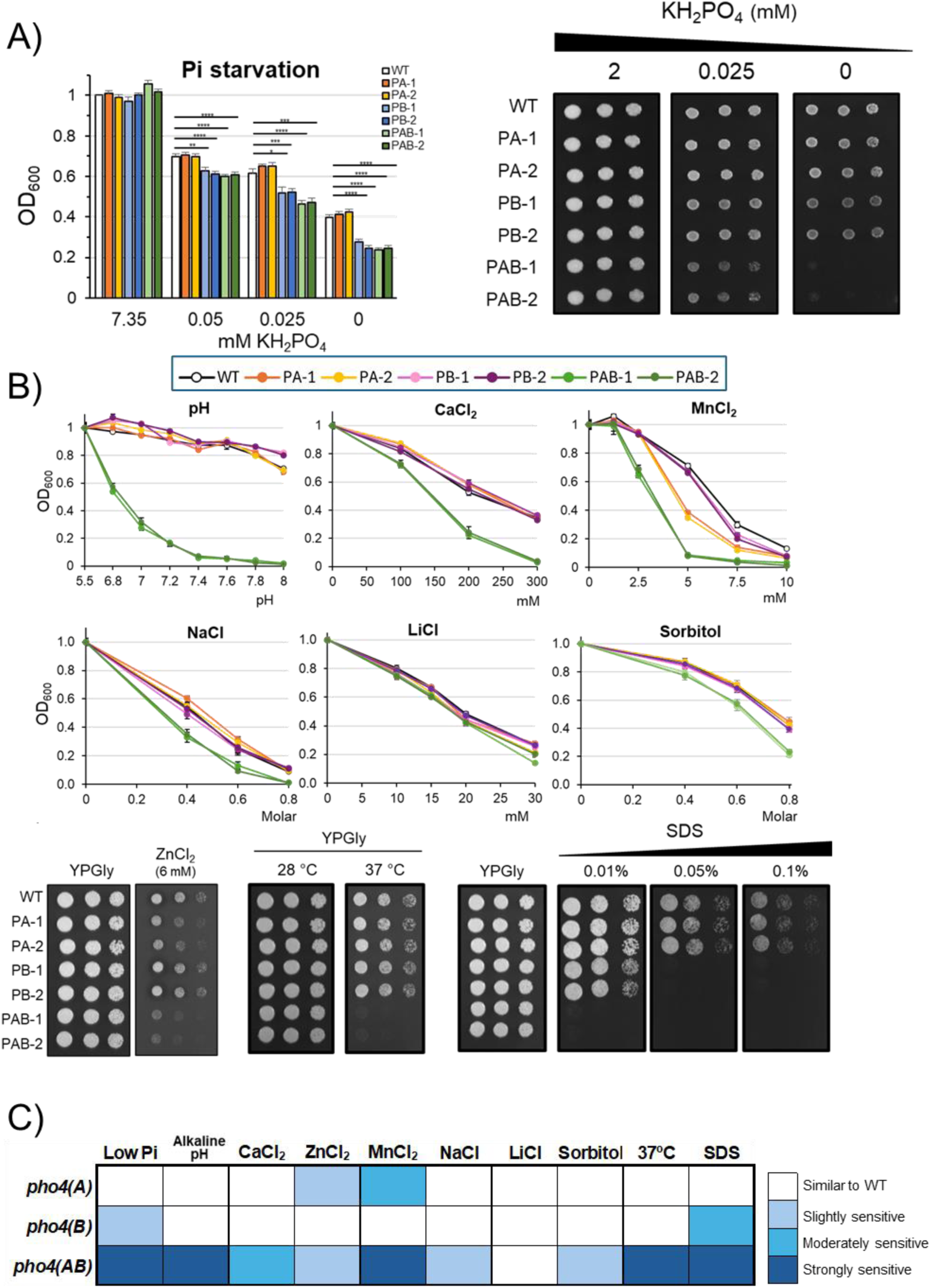
Phenotypic characterization of KpPho4(A) and KpPho4(B) single and double mutant strains. The wild-type strain X-33 (WT) and its isogenic derivatives PJA-004 (PA-1, *pho4(A)* #1), PJA-005 (PA-2, *pho4(A)* #2), PJA-021 (PB-1, *pho4(B)* #1), PJA-022 (PB-2, *pho4(B)* #2); PJA-026 (PAB-1, *pho4(A) pho4(B)* #1) and PJA-035 (PAB-2, *pho4(A) pho4(B)* #2) were grown either on synthetic medium carrying the indicated amounts of Pi (A) or (B) YPGly medium containing the indicated compounds or adjusted to the specific pHs, as described in Methods. Growth in liquid cultures was recorded after 18-20 h, and the mean ± SEM from nine independent cultures is presented. *, *p* < 0.05 / **, *p* <0.005; ***, *p* < 0.001; ****, *p* < 0.0001. Purified agar (Condalab, Cat. #1806) was used for testing growth on Pi limitation (panel A, right). Growth on plates was documented after 3 days at 28 °C. C) Summary of phenotypes detailed in panels A and B.

Altogether, these results indicate that the absence of a single transcription factor affects cell fitness only under specific conditions. Thus, lack of Pho4(B) renders cells sensitive to SDS and, importantly, to Pi deprivation, suggesting that Pho4(B) is the major component of the response to Pi starvation in *K. phaffii*. In parallel, the absence of Pho4(A) yields cells sensitive to Mn^2+^ and Zn^2+^ cations. In contrast, while concurrent lack of both transcription factors had no significant effect under standard culture conditions, it resulted in marked growth defects under nearly all stress conditions tested. This suggests that the functions of both proteins might be partially redundant and that, together, they orchestrate key mechanisms of stress resistance. It is worth noting that in all cases two different mutations were tested for each gene with virtually identical behavior. This indicates that the observed phenotypes actually derive from the absence of the encoded TF and not from unwanted off-target mutations.

### The role of Pho4(A) and Pho4(B) in the transcriptional response to Pi starvation

The influence of both *K. phaffii* Pho4 candidates on gene transcription was examined by RNA-Seq. We first considered the possible impact of the mutations in conditions of phosphate repletion (high Pi), by setting |log_2_ Fold Change| ≥1 and a *p-*value ≤0.01 as thresholds. As shown in Supplemental_Figure_S2 and Supplemental_Table_S2, lack of Pho4(A) had almost no effect, with only one gene moderately induced (*PAS_chr4_0080*, encoding an uncharacterized protein with distant relatives in other fungi). The mutation of *PHO4(B)* resulted in ten genes induced and none repressed. The induced set was highly enriched in genes encoding functionally related ATP-dependent protein folding chaperones, including Hsp60, Hsp82, Hsp78, Hsp104, Ssa1, Fes1 (Ssa1 nucleotide exchange factor), and Ssa3 (Supplemental_Fig_S2). None of these genes were found among the six induced in the double *pho4(A) pho4(B)* mutant, which included two amino acid permeases: *PAS_chr1_4_0479* (*TAT2*) and *PAS_chr4_0287* (*DIP5-1*). Ten genes were moderately repressed (2 to 3-fold) in the double mutant, including *PAS_chr4_0337* (coding for the putative high-affinity phosphate transporter Pho84-1).

Growth of wild-type cells for 24 h without phosphate resulted in a wide transcriptional response, with 803 differentially expressed genes (DEGs) of which 444 were induced and 359 repressed when compared with cells grown in the presence of Pi (Figure 3A and Supplemental_Table_S2). These values were alike for the *pho4(A)* mutant (413 genes induced and 383 repressed), but somewhat lower for the *pho4(B)* and the double mutant, mostly due to a lesser number of repressed genes (Figure 3A and Supplemental_Table_S2). A pairwise comparison of the changes occurring after Pi starvation (Figure 3B) confirmed that the mutation of *PHO4(A)* has virtually no effect on the transcriptomic profile in the absence of Pi. In contrast, lack of Pho4(B) had a noticeable impact that was amplified by the additional mutation of *PHO4(A)*.

**Figure 3.**
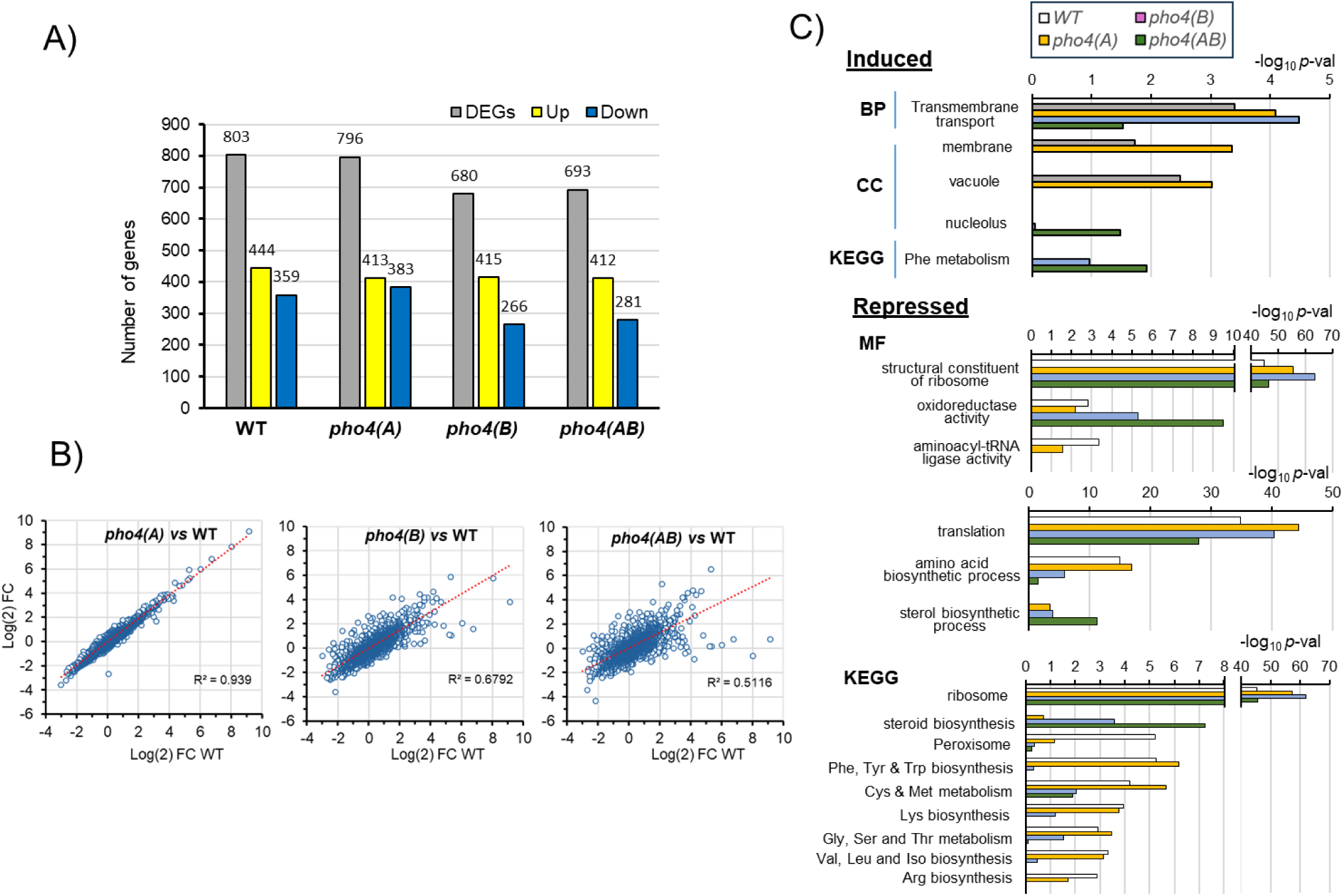
Effects of Pi starvation in the transcriptional response of wild-type and Pho4-deficient cells. A) Number of differentially expressed genes (DEGs), up-regulated or down-regulated (log2 ≥ 1 or ≤ -1 and *p*-value ≤ 0.01 ratio for the low Pi / high Pi comparison) in wild-type cells and the indicated mutants. The *pho4(A) pho4(B)* strain is denoted as *pho4(AB)*. B) Relationship between the overall transcriptional changes induced by low Pi in wild-type cells (WT) in comparison with those observed for cells mutated for *PHO4(A)*, *PHO4(B)* or both transcription factor-encoding genes. C) Gene Ontology analysis with g:Profiler of induced and repressed genes shown in panel A. MF, Molecular Function; CC, Cellular Component; BP, Biological Process; KEGG, Kyoto Encyclopedia of Genes and Genomes.

Gene ontology analysis of genes induced upon Pi starvation (Figure 3C) indicated an induction of membrane transport and vacuole-related genes that was largely abolished by lack of both Pho4 proteins. Examples were *PHO84-1* or *ENA2* transporters, as well as *VCX1*, *VPH1*, *VMA10* or *VBA1* vacuole-related genes. Phosphate starvation resulted in a drastic repression of genes involved in ribosome biogenesis and translation, and this effect was not altered by the mutations tested (Figure 3C). Genes involved in diverse amino acid biosynthetic pathways were also significantly repressed. However, in this case this effect was attenuated by mutation of *PHO4(B)* and further decreased in the double mutant, except for several genes involved in sulfur assimilation and Met and Cys biosynthesis (such as *MET3*, *MET14*, *MET17*, *MET6* or *STR3*), which were still repressed in the double mutant. Finally, lack of phosphate led to a mild but consistent decrease in the expression of the genes implicated in the biosynthesis of ergosterol from acetyl CoA (Figure 3C and Supplemental_Fig_S3), which was accentuated in the *pho4(A) pho4(B)* mutant mainly for genes involved in the second half of the pathway (biosynthesis of ergosterol from lanosterol). We then selected 28 *K. phaffii* genes corresponding to the phosphate gene regulatory network defined for *S. cerevisiae* by Ming Yip and coworkers (Ming Yip et al. 2023) and plotted their changes in expression in response to Pi starvation. As shown in Figure 4A, many well-known phosphate responsive genes were induced in the wild-type strain, including *PHO89*, *PHO5*, *VTC1*, *VTC2, VTC4,* and *GDE1*. These results fit well (Supplemental_Fig_S4) with the recent transcriptomic data obtained by Ishtuganova and coworkers (Ishtuganova et al. 2024), even though when our selection thresholds are applied, the number of DEGs identified by these authors was markedly lower (about 25% of that described in our work). We also observed induction of *PHO4(B)* itself and of upstream components of the Pi starvation signaling pathway, such as the cyclin-encoding *PHO80*, and the *PHO81* inhibitor (but not of *PHO85*). In any case, while mutation of *PHO4(A)* did not modify the responses observed for the wild-type strain, the transcriptional changes caused by Pi starvation in this set of genes was attenuated by lack of Pho4(B) and fully blocked by mutation of both TFs. It is worth noting that in *K. phaffii* there are two genes possibly encoding Pho84 high-affinity H^+^/Pi co-transporters: *PHO84-1* (*PAS_chr4_0337*) and *PHO84-2* (*PAS_chr3_0141*). Notably, *PHO84-1* was strongly induced by Pi starvation, but *PHO84-2* was repressed, a situation also observed by Ishtuganova and coworkers (Supplemental_Fig_S4). In any case, both effects were abolished in the double *pho4(A) pho4(B)* mutant. In agreement with these results, we detected a large increase in secreted phosphatase activity in wild-type cells upon phosphate starvation (Figure 4B). This increase was unaffected by mutation of *PHO4(A)* but was largely blocked by lack of Pho4(B). The vestigial activity in the latter mutant was eliminated by the additional mutation of *PHO4(A)*.

**Figure 4.**
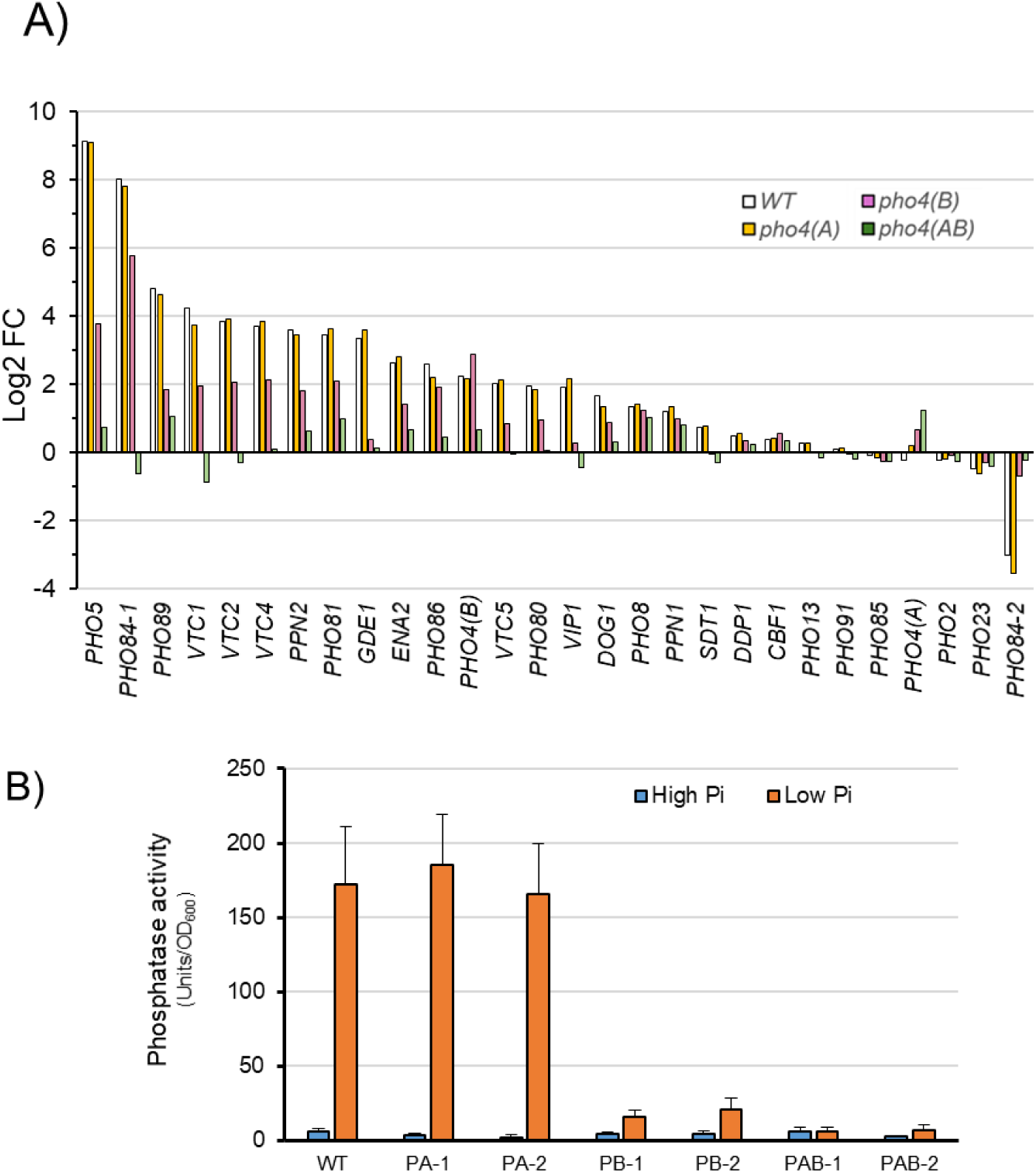
Differential effect of the *pho4(A*) and *pho4(B)* mutations on the canonical PHO regulon under Pi starvation. A) Changes in mRNA levels (low Pi vs high Pi) for members of the *PHO* regulon as defined by Ming Yip and coworkers for *S. cerevisiae* (Ming Yip et al. 2023) in wild-type cells and the indicated mutant strains. B) The indicated strains (see Fig. 2) were grown in the presence or the absence of Pi for 24 h and the activity of secreted phosphatase determined. Data are mean ± SEM from three biological samples tested in duplicate.

We then quantified (Figure 5A) the differential impact of the mutations in cells growing under Pi limitation (i.e., changes involving |log_2_ Fold Change| ≥1), and observed that no DEGs are detected when compared the *pho4(A)* and the wild-type strain, whereas 201 (96 induced and 105 repressed) were found for the *pho4(B)* strain, and 339 (157 upregulated and 182 downregulated) for the double mutant. These results confirmed that Pho4(B), but not Pho4(A), is important for the transcriptional response to Pi starvation. Gene Ontology analysis (Figure 5B) and STRING association (Supplemental_Fig_S5) revealed an increased expression of genes involved in amino acid metabolism in *pho4(B)* cells, whereas the expression of genes related to phosphate metabolism, the vacuolar ATPase, and fructose and mannose metabolism was negatively affected by the *pho4(B)* mutation. The induction of genes related to amino acid metabolism (mostly Lys, His, and branched chain and aromatic amino acids) was more intense in the *pho4(A) pho4(B)* strain (Figure 5B and 5C). Notably, three genes encoding carnitine acetyltransferases were also induced. An even wider array of genes involved in Pi homeostasis, encoding vacuolar proteins (mostly components of the vacuolar H^+^-ATPase), and genes related to ergosterol biosynthesis were found among those whose expression decreased in the double mutant strain. We then performed a computational analysis of the upstream regions of genes differentially expressed in Figure 5A in search for putative TF binding motifs. In agreement with the strong impact of the *pho4(B)* mutation on Pi homeostasis, the results revealed an enrichment in Pho4-binding consensus sequences in the genes with decreased expression in *pho4(B)* cells or the double mutant (7.6 and 5.1-fold, respectively), when compared with a random selection of genes (Figure 5D). No particular enrichment was found for other TF known to be involved in the response to other stress conditions, such as Nrg1, Crz1 or Aft1. In contrast, a moderate enrichment (1.5-fold) was detected for the Rim101 TF in genes sensitive to the lack of Pho4(B), albeit it was not found for genes sensitive to the double mutation.

**Figure 5.**
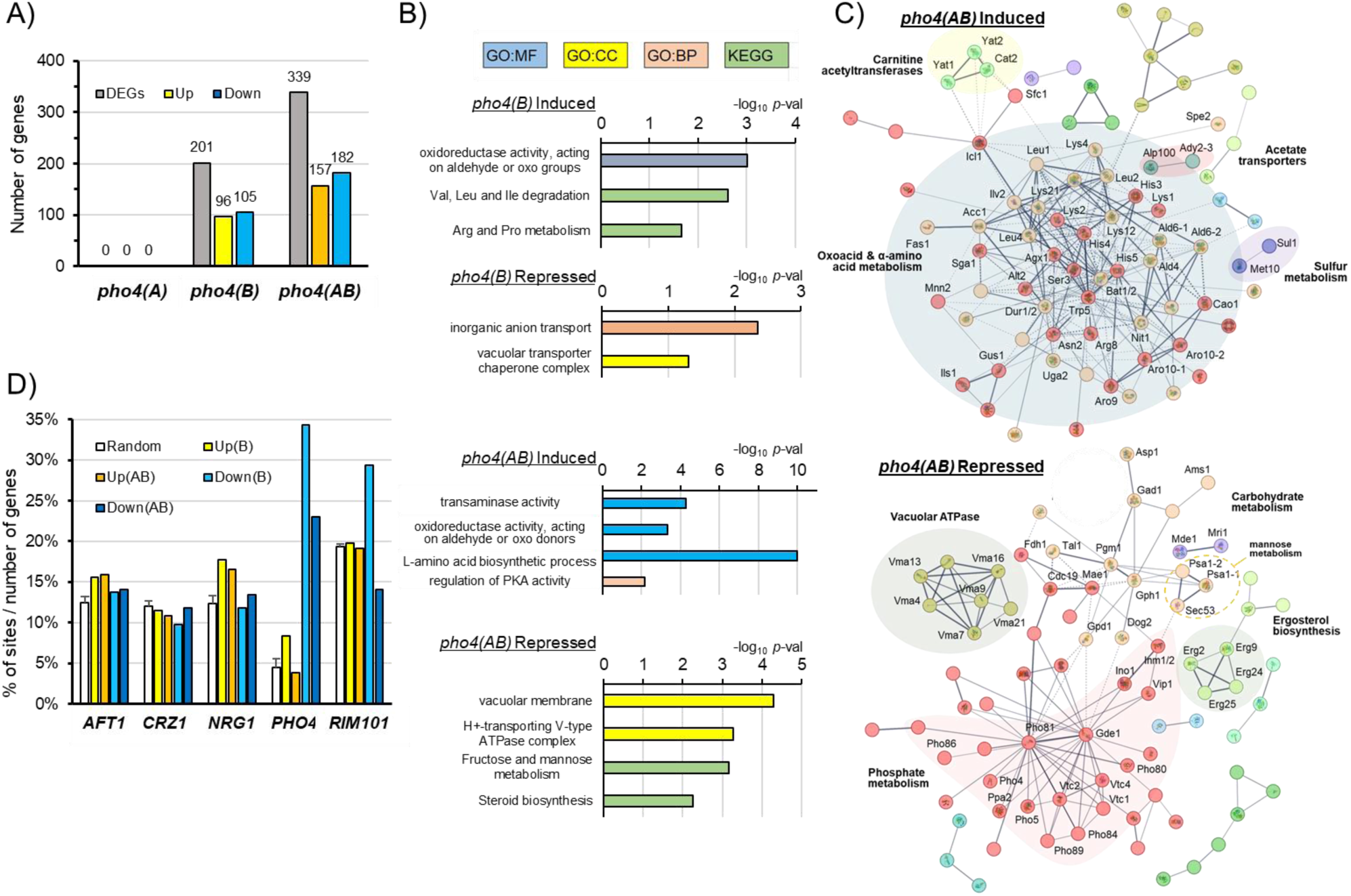
Functional analysis of genes affected by the *pho4(A)* and *pho4(B)* mutations in *K. phaffii.* A) Genes whose expression is affected by |log2 FC| ≥ 1 in the different mutants in comparison with the wild-type strain in cells growing under Pi limitation. Up, Upregulated by the mutation; Down, Downregulated. B) Gene Ontology analysis of genes induced and repressed in panel A. C) STRING network from genes differentially expressed in the double *pho4* mutant. Unconnected proteins were removed for simplicity. Colored shadows were applied manually to emphasize functional grouping. D) The upstream regions of the genes represented in panel A were analyzed for the presence of consensus sequences for the indicated transcription factors (see Supplemental Methods for details) and the number of positive hits represented as percentage over the total number of genes in each category. “Random” corresponds to the average computed from three sets of 200 randomly selected genes each.

### Pho4(A) contributes to the regulation of the expression of *PHO89* under high pH stress

We showed above that the absence of Pho4(A) had no effect on gene expression in response to Pi starvation. However, the lack of Pho4(A) aggravated the effect of the *pho4(B)* mutation not only in response to Pi shortage, but also when cells were exposed to a variety of stresses (Figure 2). We then considered the possibility that Pho4(A) might play a significant role under certain stress conditions. We tested this for the case of *PHO89*, a gene that is potently induced by both Pi shortage and alkalinization, with the aid of a reporter strain in which the *PHO89* promoter was fused to GFP. As expected (see Figure 4A), expression of GFP in response to Pi starvation was not significantly affected by lack of Pho4(A), whereas the absence of Pho4(B) yielded minimal amounts of GFP that further decreased in the double mutant (Figure 6A). Interestingly, when the same strains were subjected to alkaline stress (Figure 6B), expression of GFP at short times (30-60 min) after the onset of stress was equally decreased (≈50%) in both single mutants, while at longer times the absence of Pho4(B) became more evident. The expression of GFP was fully abolished in cells lacking both TF (Figure 6B). These results confirmed that Pho4(A) is functional in *K. phaffii* and suggest that its role might become relevant under specific stress circumstances.

**Figure 6.**
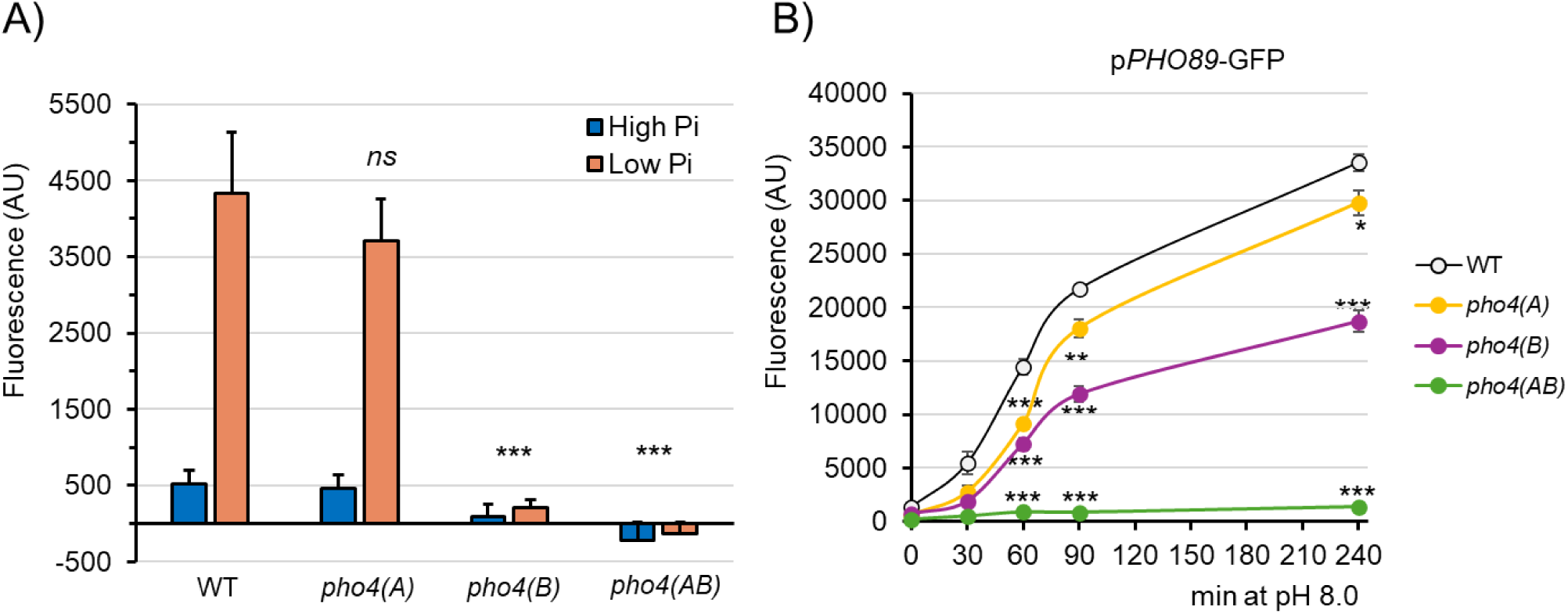
*PHO89*-driven GFP expression in response to low Pi and high pH. A) The indicated cells carrying an integrated *pPHO89-GFP* construct were grown for 24 h in medium containing 7.35 mM KH_2_PO_4_ (high Pi) or no added phosphate (low Pi). B) The indicated strains were grown to OD_600_ 0.6-0.8, and aliquots were resuspended in fresh YPGly adjusted to pH 5.5 or 8.0. Growth was resumed and samples taken at the indicated times. In all cases, samples were processed for flow cytometry as described in Methods to quantify fluorescence. Basal fluorescence produced by cells lacking the p*PHO89*-GFP insertion was subtracted. Data are mean ± SEM, n=7 (low Pi) or n=4 to 6 (high pH) independent cultures. *, *p*< 0.01; **, *p* <0.005; ***, *p* <0.0005; *ns*, not significant.

### Both *K. phaffii* Pho4 transcription factors are functional in *S. cerevisiae*

We then sought to test whether the *K. phaffii* Pho4 proteins could functionally replace Pho4 in *S. cerevisiae*. To this end, we cloned both ORFs as an N-terminally tagged 3x-HA version in a multicopy vector downstream the *ADH1* promoter and introduced them in two different genetic backgrounds (BY4741 and DBY746) widely characterized in our laboratory. However, expression of Pho4(A) from this construct in both genetic backgrounds dramatically affected cell fitness, indicating that strong overproduction of the TF was toxic for the cells (not shown). For this reason, we transferred the HA-tagged ORFs to the *tet-off* pCM189 vector, which allows centromeric expression only in the absence of doxycycline. The vectors bearing native *K. phaffii PHO4(A)* and *PHO4(B)* ORFs, as well as their mutated versions *pho4(A)*Δ110 and *pho4(B)*Δ116 (obtained from strains PJA-036), were introduced into the above-mentioned strains and their *pho4*Δ derivatives. We observed (Figure 7A) that in the absence of doxycycline the expression of both native KpPho4 isoforms (but not that of their mutated versions) in the DBY746 *pho4*Δ strain allowed growth with similar efficiency in medium lacking added Pi. When these cells were grown at alkaline pH, KpPho4(B) was able to confer normal tolerance, but KpPho4(A) provided only marginal growth. Surprisingly, when the same plasmids were introduced in strain BY4741 (Supplemental_Fig_S6A), only the native version of Pho4(A) improved growth on plates lacking added Pi or at pH 8.0, whereas expression of Pho4(B) provided marginal growth only at alkaline pH. When we checked the levels of KpPho4(A) and KpPho4(B) in both strains we found that in the DBY746 background (Figure 7B) the amounts of KpPho4(A) and KpPho4(B) were quite similar, whereas in BY4741 the level of KpPho4(B) was substantially lower than that of KpPho4(A) (Supplemental_Fig_S6B). We considered the possibility that the failure of KpPho4(B) to complement the *pho4* mutation in the BY4741 background could be due to insufficient levels of the transcription factor.

**Figure 7.**
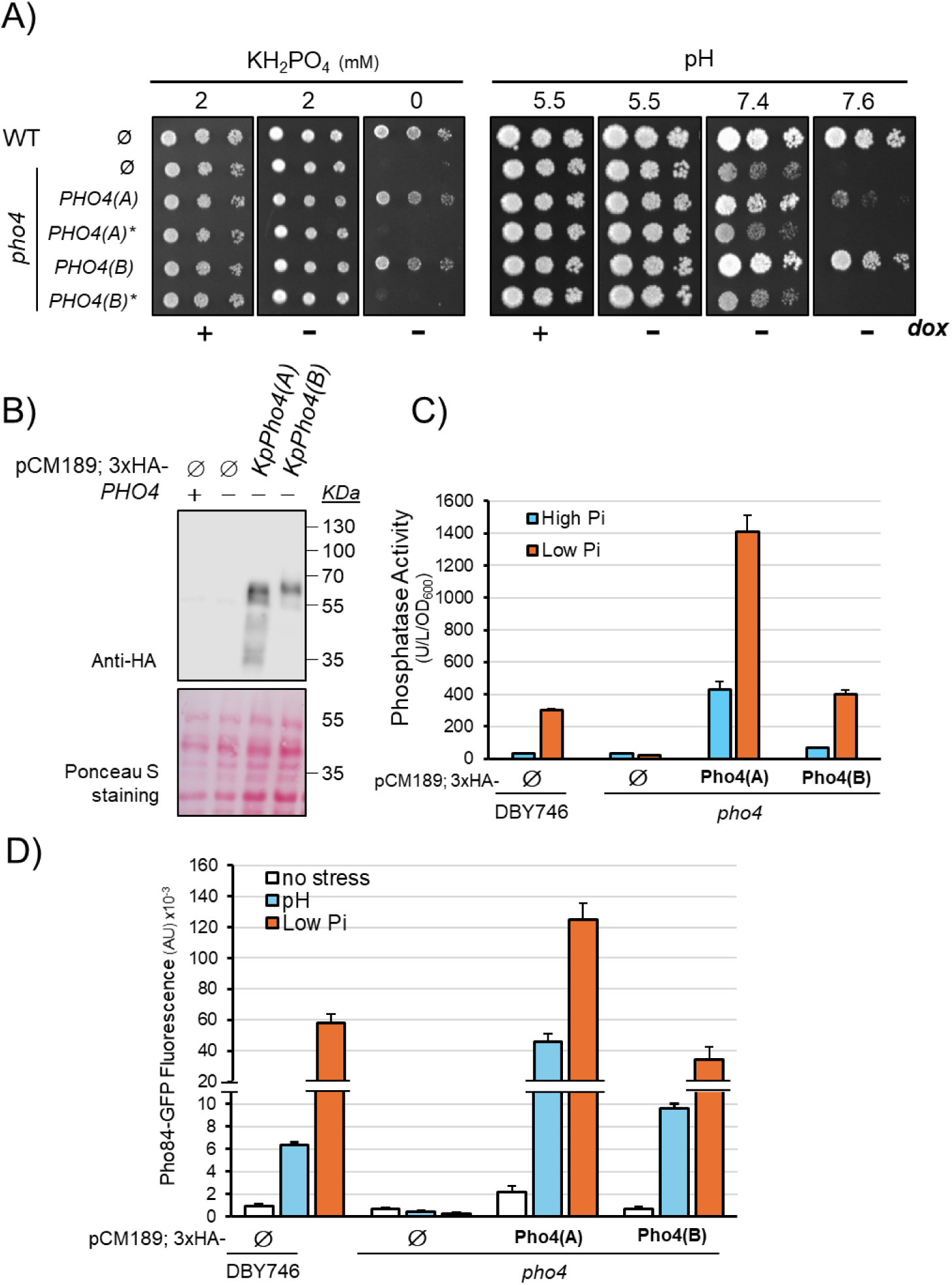
The expression of Pho4(A) and Pho4(B) complements the *pho4* mutation in *S. cerevisiae*. A) Strain DBY746 and its derivative MAC008 (*pho4::kanMX*) were transformed with the empty vector pCM189 (Ø) or the same vector carrying 3xHA-tagged version of *KpPHO4(A)*, *KpPHO4(B)*, or their mutated versions (denoted by an asterisk). Transformants were spotted on synthetic medium (Pi free) plates at pH 5.5 carrying the indicated amounts of Pi or on YPD at the indicated pH (adjusted with 50 mM TAPS), and growth recorded after 6 or 4 days at 28 °C, respectively. B) The mentioned strains were transformed as above. Samples were taken after 20 h of growth in the absence of doxycycline to allow expression of the heterologous Pho4 proteins, subjected to SDS-PAGE (10 µg protein), transferred to membranes and probed with anti-HA antibodies. The Ponceau stained membrane is shown for loading comparison. C) Strains DBY746 and MAC008 (*pho4*) were transformed as in panel B. After 20 h of growth in the absence of doxycycline, cells were washed and resuspended in Pi-free medium (low Pi) or with added 7.35 mM KH_2_PO_4_ (high Pi), and growth resumed for 6 h prior determination of phosphatase activity. Data are mean ± SEM from four biological replicas tested in duplicate. D) DBY746 and MAC008 (*pho4*) cells were co-transformed with the indicated pCM189-based vectors plus YCp111-*PHO84*-*GFP*. Cultures were treated as described in Methods and processed for flow cytometry measurement. Samples were taken after 2 h at pH 8 (pH) or after 6 h upon Pi starvation (low Pi). Data are mean ± SEM from four biological replicates.

To test this possibility we monitored the activity of secreted phosphatase, a known target for Pho4 in *S. cerevisiae*, in response to Pi starvation in both wild-type and Pho4-deficient backgrounds. In the wild-type DBY746 background (Figure 7C) phosphatase activity increased about 9-fold upon Pi starvation. This value was very similar to that obtained by expression of KpPho4(B) in the *pho4* background. Expression of KpPho4(A) in the *pho4* mutant (MAC008 strain) led to significant levels of phosphatase activity even under repressive conditions (high Pi) and the activity under Pi starvation exceeded nearly five-fold that of wild-type cells. In contrast, in the BY4741 *pho4* background KpPho4(A) was able to produce about 65% of the level generated by the native ScPho4 transcription factor, but cells expressing KpPho4(B) did not show significant enzyme activity (Supplemental_Fig_S6C).

We finally tested the induction of the *S. cerevisiae* high-affinity H^+^/Pi transporter Pho84 (whose expression is needed for growing under Pi limitation) by expressing from the *PHO84* promoter a C-terminally GFP tagged version of Pho84 and monitoring the produced fluorescence both in the Pho4-deficient DBY746 and BY4741 backgrounds. As shown in Figure 7D, expression of Pho84 increased about six-fold in response to alkalinization and nearly 60-fold upon Pi starvation in DBY746 cells, and this response was fully abolished in the MAC008 mutant (*pho4*Δ). Expression of KpPho4(B) in strain MAC008 restored Pho84 expression to nearly wild-type levels under both stresses, whereas Pho84 levels upon expression of KpPho4(A) exceeded in seven-fold (alkaline pH) and two-fold (Pi starvation) those observed for the wild-type strain. Interestingly, the level of Pho84 in the Pho4-deficient BY4741 background in response to Pi limitation when KpPho4(A) was expressed (Supplemental_Fig_S6B) was largely above that obtained in the wild-type but almost undetectable and far below those derived from native Pho4 when KpPho4(B) was tested.

Overall, these results indicate that both *K. phaffii* Pho4 TFs are able to substitute for *S. cerevisiae* Pho4 in the response to Pi starvation in the DBY746 background. The inability for KpPho4(B) to do so in the BY4741 background can be explained by the failure to properly induce the Pho84 transporter and the secreted phosphatase, possibly because of insufficient levels of the TF.

## DISCUSSION

While duplication or expansion of downstream phosphate transporter or phosphatase gene families regulated by Pho4 is not unusual in fungi, it is widely accepted that Pho4-encoding genes exist as a single instance (Gomes-Vieira et al. 2018; Tomar and Sinha 2014). However, we show here that *K. phaffii* contains two genes encoding relatively distant proteins that function as Pho4 transcription factors. In fact, we have detected the presence of two likely Pho4 proteins only in the proteome of the very close relative *Komagataella kurtzmanii* (LJB42_003486 and LJB42_000190). In contrast, the related species *Brettanomyces naardenensis*, *Pichia inconspicua*, and *Pichia kluyveri* show a single protein that could reasonably qualify as Pho4. Therefore, the situation in *K. phaffii* is exceptional and, given the relevance of this organism as host for foreign protein expression, it was worth examining it in detail.

Our transcriptomic profiling of cells deprived of Pi for 24 h yielded 811 DEGs. This number is almost 4-fold higher (when our threshold is applied) than that recently reported by Ishtuganova and coworkers (Ishtuganova et al. 2024). Because in both cases the X-33 strain was employed, the composition of the medium was virtually identical, and the starvation period was the same, we assume that a difference in the quantitative coverage of reads or the homogeneity among replicates might explain the higher number of DEGs reported here. In any case, we could confirm the induction of many genes required for efficient Pi transport or scavenging (Figure 4A and S4), as well as the opposite behavior of the two Pho84-encoding genes under Pi starvation. We also detected repression of genes involved in the metabolisms of specific amino acids, such as Met and Cys, as well as a widespread repression of genes encoding ribosomal proteins and translation factors. In contrast, we did not find significant alterations in genes related to glycolysis/gluconeogenesis or the fatty acid β-oxidation pathway reported by these authors (Ishtuganova et al. 2024).

Our results show that Pho4(B) has a major role in the response to Pi starvation, while the single deletion of *PHO4(A)* has virtually no effect on transcription (Figs. 3-5). Interestingly, lack of both TF often results in full elimination of the response, suggesting that, in the absence of Pho4(B), Pho4(A) gains some relevance. In fact, although *PHO4(A)* is not induced by Pi starvation, its mRNA levels increase modestly but significantly in the *pho4(B)* mutant (Figure 4A and Table S3), and even more in the double mutant. Thus, in the absence of Pho4(B) an increase in *PHO4(A)* expression might contribute to sustaining part of the response to Pi shortage and would explain why, despite its prominent role in the transcriptional response, deletion of *PHO4(B)* has a partial effect on growth under Pi limitation (Figure 2A) and the elimination of both TF is needed to fully block growth. In contrast, the expression of *PHO4(B)* is induced under Pi shortage and it is largely blocked in the absence of both TF (Figure 4A and Supplemental_Table_S2). This could be justified by the presence of a possible Pho4-binding site at position -241 (CCACGTGG, *p*-val 3.5E-05). Notably, lack of phosphate does not alter Pho4 levels in *S. cerevisiae* or *C. glabrata* (Kerwin and Wykoff 2009), but induction of the *CnPHO4* gene under Pi limitation has been reported for *Cryptococcus neoformans* (Toh-e et al. 2015). Thus, some fungi may resort to self-regulation of *PHO4* expression to allow rapid adaptation to Pi starvation.

Pho4(B) is required not only for the induction of genes involved in phosphate transport and metabolism (the canonical *PHO* regulon) but also affects other aspects of yeast physiology (Figure 5). For instance, lack of Pi triggers a modest but consistent repression of many genes involved in ergosterol biosynthesis and the repressive effect is accentuated by mutation of *PHO4(B)* and further by lack of both TF (Supplemental_Fig_S3). We are not aware of this effect being described for other yeast. However, previous results (Albacar et al. 2025, 2023) showed the repression of several of these genes (*ERG1*, *ERG25*, and *ERG3*) under alkaline pH. Notably, similar changes upon exposure to alkaline pH have been documented in *A. nidulans* (Picazo et al. 2020) and *C. albicans* (Garnaud et al. 2018). These similarities reinforce the notion that Pi shortage and alkaline stress are closely related.

In accordance with (Ishtuganova et al. 2024) we also observe that lack of phosphate results in *K. phaffii* in widespread repression of genes relevant to ribosome biogenesis (including ribosomal proteins) as well as of translation factors. This reaction, as well as the repression of ergosterol biosynthetic genes, points to a halt in proliferation. A strong repression upon Pi starvation of genes associated with the ribosome and translation was also described in *S. pombe* (Garg et al. 2023), and it was partially relieved by mutation of *PHO7* (Garg et al. 2024). A similar pattern was described in *N. crassa*, although in this case it was shown that the effect was directly mediated by NUC-1, the Pi-sensitive TF that also mediates most of the response to Pi starvation in this organism (Huberman et al. 2025). In contrast, our data shows that in *K. phaffii* repression of ribosome and translation-related genes does not depend on any of the transcription factors (Figure 3C and Supplemental_Table_S2).It should be noted that this effect on ribosomal-related gene repression is not general for all fungi, as it has not been reported for *S. cerevisiae*, *C. neoformans* (Toh-e et al. 2015) or *C. albicans* (Ikeh et al. 2016).

Our data show that, collectively, Pho4(A) and Pho4(B) regulate over 300 genes in response to Pi starvation, of which 182 were less expressed in the absence of both TFs (Figure 5A). This far exceeds the 20-30 Pho4-regulated genes reported for *S. cerevisiae* (Ogawa et al. 2000; Ming Yip et al. 2023), but it is in the range of the nearly 80 genes requiring Pho4 for induction by Pi starvation in *C. glabrata* (He et al. 2017), or the around 150 genes identified in *C. albicans* (Ikeh et al. 2016) or *C. neoformans* (Toh-e et al. 2015). Interestingly, this expansion in the range of Pho4-regulated genes has been linked to a lower or null functional dependence for the association with the Pho2 co-activator and a gain of additional regulatory functions (Köhler et al. 2020; He et al. 2017; Lev and Djordjevic 2018). Among these additional functions, Pho4 was revealed as a key element in the expression of stress response genes. Thus, a *C. albicans pho4* null mutant was sensitive not only to lack of phosphate, but also to sorbitol, NaCl, MnCl_2_, CaCl_2_, H_2_O_2_ (peroxide stress), and alkaline stress (pH 8), among other stresses (Urrialde et al. 2016; Ikeh et al. 2016). Lack of the Pho4 TF in *C. neoformans* rendered cells sensitive to calcofluor white, amphotericin B, CaCl_2_, and nitrosative stress, although this was observed only under Pi limitation, whereas these cells were sensitive to alkaline pH even in the presence of normal Pi levels (Lev et al. 2017). *K. phaffii* contains a possible Pho2 TF-encoding gene (*PAS_chr2_2_0043*) (Figure 1A), albeit its level of identity with Pho2 from *S. cerevisiae* is rather low (28%). It would be interesting to explore in the future whether it is required for any of the Pho4 isoforms to function.

In concordance with above, we show here that *K. phaffii* cells lacking Pho4(A) and Pho4(B) are not only sensitive to lack of Pi in the medium, but also strongly sensitive to different kinds of stress. Interestingly, single mutants display different specific phenotypes. Thus*, pho4(A)* cells are sensitive to high levels of Mn^2+^ and Zn^2+^, whereas *pho4(B)* is sensitive to SDS (Figure 2), and in all cases the phenotype of the single mutant is aggravated by the double mutation. We have observed that expression of the *ENA2* ATPase, a key element for the tolerance to saline and alkaline stress (Haro et al. 1991; Platara et al. 2006; Ramos et al. 2011) is induced by Pi starvation and that such induction is virtually abolished in the double mutant (Table S3). Since the double mutant is strongly sensitive to alkaline pH, this might suggest that Pho4 TFs could be also involved in the upregulation of *ENA2* upon alkaline stress. Because of the functional relationship between Pi starvation and alkaline stress responses, it would be most interesting to explore the role of Pho4 in adaptation to alkaline stress at a genome-wide level. In contrast, the observation that the double mutant was sensitive to NaCl and sorbitol but not to the more toxic analog LiCl (Figure 2B), argues against an involvement of Pho4 in the response to cation toxicity and points to a role on osmotic homeostasis. It must be noted that mutation of Pho4 also causes osmotic sensitivity in *C. albicans* (Urrialde et al., 2016). Notably, osmotic defects have been historically associated to alterations in the cell wall integrity (CWI) pathway based on the Slt2 MAP kinase cascade and its crosstalk with accessory pathways, such as TOR, calcineurin or the high-osmolarity pathway (Fuchs and Mylonakis 2009), but no mechanistic link with the PHO pathway has been reported so far. In this regard, we found that three genes encoding carnitine acyl transferases (*YAT1*, *YAT2* and *CAT2*) are found induced upon Pi starvation in the double mutant, but not in wild-type cells (Figure 5C and Table S3). These genes have been reported to be relevant for adaptation to diverse forms of stress in S*. cerevisiae* (Franken et al. 2008). Therefore, the induction observed here could be interpreted as an attempt to reinforce their stress response arsenal weakened by lack of both Pho4-encoding genes. In any case, our transcriptomic profiling cannot provide an explanation for some of the observed phenotypes. It is particularly intriguing the sensitivity to Mn^2+^ ions of the *pho4(A)* and *pho4(A) pho4(B)* strains, since we do not observe in these mutants changes in expression of the SFM transporters known to mediate Mn^2+^ transport (Cohen et al. 2000). A mild effect of the *pho4* mutation on Mn^+2^ sensitivity has been described for the *pho4* mutant in *C. albicans* (Ikeh et al. 2016), and it was linked to the inability of this strain to store polyphosphate. However, given the lack of effect of the *pho4(A)* mutation alone on Pi homeostasis, such explanation does not seem likely in our case.

We show here that expression of both *K. phaffii* Pho4 isoforms can complement the *pho4* mutation when expressed in *S. cerevisiae*. This implies that both *K. phaffii* TFs are responsive to the *S. cerevisiae* Pi-sensitive regulatory network, as deduced from the increase of expression of the Pho84 transporter and the induction of secreted phosphatase activity in response to Pi starvation (Figures 7C,D and S6B,C). However, the higher-than-normal phosphatase activity and Pho84 levels in the presence of Pi when expressing Pho4(A) in both DBY746 and BY4741 backgrounds suggest that this TF might escape somewhat from the inactivation mechanisms. In any case, KpPho4(A) appears not only fully functional in triggering Pi-starvation related responses in *S. cerevisiae*, but even more efficient than KpPho4(B) (Figs. 7C and D).This is somewhat surprising, because our data show that in *K. phaffii* KpPho4(B) is the major contributor in controlling expression of genes related to phosphate homeostasis. It could be proposed that the consensus sequence for Pho4 binding could be somewhat different in *K. phaffii* vs *S. cerevisiae*. However, prediction of putative Pho4 binding sites in the promoter for 9 phosphate-related, Pho4(B)-regulated genes, in comparison to their homologs in *S. cerevisiae* yielded a very similar pattern (Supplemental_Fig_S6D). A differential feature between KpPho4(A) and KpPho4(B) is that, in the latter, the Ser at phosphorylation site 6 reported in *S. cerevisiae* is missing (Figure 1B). Remarkably, it has been shown that mutation of this Ser to Ala reduced the ability of the TF to activate transcription of acid phosphatase (Komeili and O’Shea 1999). Therefore, the absence of this phosphorylatable residue might explain the limited efficiency of KpPho4(B) in comparison with KpPho4(A) when expressed in *S. cerevisiae*.

In conclusion, we demonstrate here an exceptional genetic feature among yeasts: the presence in *K. phaffii* of two functional Pho4-encoding genes. In combination, their protein products play a key role not only in maintenance of phosphate homeostasis, but also in stress response. Since Pho4-regulated genes, such as *PHO89*, have been successfully used to drive expression of industrially relevant proteins both under Pi limitation (Ahn et al. 2009; Xie et al. 2020) or alkalinization of the medium (Albacar et al. 2024), gaining knowledge about how Pho4 TFs are regulated will help to set the basis for the development of improved protein expression platforms.

## METHODS

### Growth conditions and recombinant DNA techniques

The DH5α *Escherichia coli* strain was grown at 37 °C in LB medium. When used for plasmid host, the medium was supplemented with hygromycin B (100 µg/ml). *E. coli* transformation and standard recombinant DNA techniques were performed as described (Green and Sambrook 2012). *K. phaffii* cells (X-33) were grown at 28 °C on YP medium (Yeast extract 1%, Peptone 2%) supplemented with 2% glycerol (YPGly) as carbon source (unless otherwise stated) and were transformed by electroporation as described in (Lin-Cereghino et al. 2005) with a Gene Pulser electroporator (Bio-Rad). *S. cerevisiae* cells used in this work are BY4741 (*MAT**a** his3*Δ1 *leu2*Δ0 *met15*Δ0 *ura3*Δ0) and its isogenic *pho4::kanMX4* derivative (Giaever et al. 2002; Brachmann et al. 1998), and DBY746 (*MAT*α *his3*Δ*1 leu2-3 leu2-112 ura3-52 trp1-289*) and its isogenic derivative MAC008 (*pho4::kanMX*). Strain MAC008 was prepared by transformation of DBY746 cells with a *pho4::kanMX4* cassette amplified with oligonucleotides pho4_5’_amp and pho4_3’_amp (Supplemental_Table_S3A) from the genomic DNA of strain ASC15 (Serra-Cardona et al. 2014). *S. cerevisiae* cells were grown at 28 °C in YP medium supplemented with 2% glucose (YPD), or in synthetic medium (SC) lacking uracil and leucine when required (Adams et al. 1998) and were transformed by a variant of the lithium acetate method (Gietz et al. 1995). For growth on plates, 2% of bacteriological agar (Condalab) was added to the medium, which was replaced by purified agar when testing for growth under Pi limitation (Condalab, Cat. #1806).

### Generation of *K. phaffii PHO4(A)* and *PHO4(B)* mutants

Different mutations were introduced in the ORFs of *PHO4(A)* (*PAS_chr1-1_0265*) and *PHO4(B)* (*PAS_chr2-1_0177*) by CRISPR/Cas9-mediated gene disruption (Gassler et al. 2019). To this end, two different single-guide RNAs (sgRNAs) were constructed for each gene (Supplemental_Table_S3B) by overlap extension PCR using oligonucleotides described in Supplemental_Table_S3A and Q5 DNA polymerase (New England Biolabs). The fragments were purified by agarose gel electrophoresis and cloned by a Golden Gate assembly reaction into the vector BB3cH_pGAP_23*_pLAT1_Cas9 (Gassler et al. 2019; Prielhofer et al. 2017) taking profit of the flanking BpiI restriction sites. This vector expresses hCas9 from the *LAT1* promoter and can be selected in the presence of hygromycin (Addgene # 104907). The CHOPCHOP v3 web toolbox (Labun et al. 2019) was employed for selection of the Cas9 target sequences. *K. phaffii* strain X-33 was transformed with the different constructs and plated on YPD agar containing hygromycin (200 µg/ml). Colonies were processed as in (Albacar et al. 2025) and the mutations identified by sequencing the relevant genomic region with oligonucleotides listed in (Supplemental_Table_S3A). Double mutations were made by transformation of strain PJA-004 (*pho4(A)*Δ110) with constructs p907_sgRNA-Pho4(B).1 and p907_sgRNA-Pho4(B).2 (Supplemental_Table_S3B) and selected as above. The specific genotype of these strains and the effect of the mutations on the produced protein is shown in Supplemental_Table_S4).

For evaluation of GFP expression from the *PHO89* promoter, the p*PHO89*-GFP translational reporter described in (Albacar et al. 2023) was digested with PmeI (which cuts once within the *AOX1* promoter) and 100 ng of DNA were used to transform the appropriate strains (Supplemental_Table_S4). Transformants were recovered in YPD agar plates containing hygromycin (200 µg/ml). The correct insertion of the plasmid into the chromosomal *AOX1* locus was confirmed by PCR. The wild-type strain carrying the p*PHO89*-GFP insertion (PJA-037) was previously described (Albacar et al. 2023).

### RNA preparation and transcriptomic analysis

For assessment of the transcriptional response of *K. phaffii* to phosphate starvation, cells were grown for ∼40 h on 20 ml of synthetic medium prepared with Formedium premade mix (ref. CYN0803, Yeast Nitrogen Base without Amino Acids and Phosphate) supplemented with 15 mM KCl, 2% glycerol and 7.35 mM KH_2_PO_4_, and adjusted to pH 5.8. For cultures to be subjected to Pi starvation (low Pi), cells (10 OD_600_) were centrifuged and washed twice with medium without KH_2_PO_4_, whereas for control cultures (high Pi) 2.5 OD_600_ were taken and washed twice with medium containing 7.35 mM KH_2_PO_4_. In all cases, cells were resuspended in 50 ml of the appropriate medium and growth resumed for 24 h. Then, ∼5 OD_600_ of each culture was collected, placed on ice, and immediately centrifuged (5 min at 1000 xg) at 4 °C. Then, one ml of cold H_2_O was added, and cells were transferred to pre-cooled 1.5 ml screw cap tubes and centrifuged (two min at 1000 xg) at 4 °C. Finally, the supernatant was removed, and samples were snap-frozen in liquid N_2_ and stored at -80 °C.

Total RNA extraction was carried out as described in (Albacar et al. 2023) from three independent cultures of wild-type, *pho4(A)* (strain PJA-004), *pho4(B)* (strain PJA-021) and *pho4(A) pho4(B)* (PJA-026) cells. RNA concentration and integrity were assessed by Qubit assay (ThermoFisher) and with an Agilent 5400 Fragment Analyzer System, respectively. Messenger RNA was purified from total RNA using poly-T oligo-attached magnetic beads. After fragmentation, synthesis of the first strand cDNA was achieved by means of random hexamer primers, followed by the second strand cDNA synthesis using dTTP. After end repair, A-tailing, and adapter ligation, fragments were size selected, PCR amplified, and purified. Libraries were then pooled and subjected to sequencing in a NovaSeq X Plus equipment (Illumina) by Novogene Co., Ltd.

Raw reads (fastq format) were transformed into clean reads through fastp (Chen et al. 2018). An average ± SD of 2.4E7 ± 3.3E6 clean reads (3.62± 0.49 clean Gbases) were obtained and 97.2% ± 0.18 of them reached Q30 or above. The HISAT2 (v2.2.1) software (Kim et al. 2019) was used to align paired-end clean reads to the *K. phaffii* GS115 reference genome (assembly GCA_000027005.1), resulting in an average of 95.02% ± 1.78 of mapped reads. Differential gene expression analysis was carried out using DESeq2 (Love et al. 2014) by establishing a |log2(Fold Change)| >= 1 and a *p*-value <= 0.01.

Gene Ontology enrichment analysis was performed with g:Profiler (Kolberg et al. 2023) using the *K. phaffii* GS115 data from the Ensemble Genomes fungi database and the default settings. Interaction networks between proteins were created with the STRING database v12.0 (Szklarczyk et al. 2023) with a Minimum Interaction Score of 0.5 (medium-high confidence) and K-means clustering (number of clusters ranged from 6 to 10 depending on the protein set used).

### Phenotypic assays

*K. phaffii* and *S. cerevisiae* strains were assessed for diverse phenotypes in both liquid and solid media. For studies in liquid medium, exponential cultures were inoculated at OD_600_ = 0.004 in 300 µl of the appropriate medium in Bioscreen honeycomb microplate plates (#9502550) and cultured for 18-24 h 28 °C in an Ika-Schuttler MTS-2 orbital shaker at 900 strokes/min. To evaluate the ability to grow under phosphate limitations, the Formedium premade mix (ref. CYN0803) was used as described in the transcriptomics section. For phenotyping in solid media, cultures were diluted up to at OD_600_ = 0.05, and 5-fold dilutions were prepared. Three µl of the dilution were spotted on the appropriate plates and grown for the indicated times prior documentation. For low Pi plates, 1.5% of purified agar (Condalab, Ref. 1806) was used to ensure a Pi concentration < 15 µM.

### Expression of GFP from the *PHO89* promoter in *K. phaffii*

To monitor the response to phosphate starvation and to alkalinization, the appropriate strains carrying the p*PHO89*-GFP insertion were grown to saturation on 5 ml of Pi-deficient medium supplemented with 15 mM KCl plus 2% glycerol and 7.35 mM KH_2_PO_4_, adjusted to pH 5.8. The cultures were then split in two 2.5 ml aliquots and centrifuged (5 min, 1000 xg) at room temperature. One aliquot was washed twice with the same medium lacking the phosphate salt (Low Pi) and resuspended with 2.5 ml of the same medium. The other aliquot was treated as above but using medium supplemented with 7.35 mM KH_2_PO_4_ (High Pi). After resuspension, cultures were incubated at 28 °C with shaking for 24 hours. To evaluate the response to alkalinization, o/n cultures on YPGly were diluted to OD_600_ = 0.3, split into two 5-ml aliquots and growth resumed until OD_600_ = 0.6-0.8. Then one aliquot was resuspended in 5-ml fresh YPGly with 50 mM TAPS and adjusted to pH 5.5 and the other in the same volume of YPGly with 50 mM TAPS adjusted to pH 8.0 and growth was resumed. In all cases, 875 µl-samples were collected at different times and processed for flow cytometry as in (Albacar et al. 2023). The fluorescence of the samples was analyzed in a CytoFLEX (Beckman Coulter) flow cytometer using the FIT-C filter (525/40).

### Determination of Pho84-GFP expression by flow-cytometry

Wild-type DBY746 and its *pho4*Δ derivative MAC008 were co-transformed with vector YCp111-PHO84-GFP together with either plasmids pCM189 (empty vector), pCM189-KpPHO4(A) or pCM189-KpPHO4(B). Cells grew overnight until saturation in synthetic media lacking uracil and leucine supplemented with 100 µg/ml doxycycline. One OD_600_ of cells was collected, centrifuged, washed with the same medium without doxycycline and resuspended in 10 ml of this medium. After growth for 20 h in the absence of doxycycline an aliquot (875 µl) was collected as a control sample and fixed as described above. From the remaining volume of the culture, two aliquots of 3 ml were taken. One aliquot was washed twice with low phosphate medium supplemented with 7.35 mM KH_2_PO_4_ at pH 8.0, resuspended in 3 ml of the same media and growth resumed for 2 hours (high pH). After this time, an 875 µl sample was taken and fixed. The other 3 ml aliquot underwent the same procedure with media lacking added Pi and pH adjusted to 5.8. In this case samples were fixed after 6 hours of growth. Samples were analyzed in a CytoFLEX equipment (Beckman Coulter) as described above.

### Determination of secreted phosphatase

To determine secreted acid phosphatase activity in response to phosphate starvation in *K. phaffii*, wild-type X-33 cells and *pho4(A)*, *pho4(B)* and *pho4(A) pho4(B)* derivatives (two different variants each) were treated as described for transcriptomic analysis. In this case, however, only 0.5 and 2 OD_600_ were collected for subsequent growth on high or low Pi, respectively, and in both cases, cells were resuspended in 10 ml of fresh medium after appropriate washing. Growth was resumed and samples of one ml were taken at the indicated times, washed with cold MilliQ water, and phosphatase activity was measured in a suspension of intact cells essentially a described in (Payne et al. 1995) except that incubation was performed for 30 min and released *p*-nitrophenol was measured at 405 nm. For determination of phosphatase activity in *S. cerevisiae*, the relevant strains transformed with pCM189-derived constructs were grown to saturation in synthetic medium lacking uracil in the presence of doxycycline. The appropriate volume of culture to yield an OD_600_ of 0.05 when resuspended in a final volume of 10 ml was centrifuged and washed with the same medium lacking doxycycline. Growth continued for 20 h and then two one-ml aliquots were taken. One aliquot was washed twice with Pi-free medium containing 7.35 mM KH_2_PO_4_ and finally resuspended in one ml of the same medium, while the other was processed equally but using medium lacking added Pi. Growth was resumed for 6 h and cells were processed as above for determination of phosphatase activity.

Construction of vectors expressing KpPho4(A), KpPho4(B) and Pho84-GFP in *S. cerevisiae,* immunoblotting, and diverse computational analyses are described in Supplemental Methods.

## DATA ACCESS

All raw and processed sequencing data generated in this study was deposited at NCBI’s Gene Expression Omnibus (GEO; https://www.ncbi.nlm.nih.gov/geo/) under study GSE310523.

## ACKNOWLEDGEMENTS

We thank Montserrat Robledo for skillful technical support. The efficient support of SCAC and SGiEB UAB facilities is acknowledged. We greatly appreciate the efforts of Bruno Contreras Moreira (EEAD-CSIC, Spain) in making available *K. phaffii* analysis at the plant RSAT server. Work supported by grants PID2020-113319RB-I00 and PID2023-150535OB-I00 (AEI, Ministerio de Ciencia, Innovación y Universidades) to JA and AC, and 2023 PROD 00006 (AGAUR, Generalitat de Catalunya) to JA. RW is a recipient of a China Scholarship Council (CSC) fellowship.

## AUTHOR CONTRIBUTIONS

M. Albacar: Investigation; Formal analysis; Writing - review & editing. A. González: Investigation; Formal analysis; Writing - review & editing. R. Wang: Investigation; A. Casamayor: Investigation; Formal analysis; Funding acquisition; Writing - review & editing. J. Ariño: Investigation; Conceptualization; Formal analysis; Funding acquisition; Supervision; Project administration; Writing - original draft; Writing - review & editing.

